# A comprehensive reference database to support untargeted metabolomics in *Pseudomonas putida*

**DOI:** 10.64898/2026.03.20.713193

**Authors:** Dylan H. Ross, Christine H. Chang, Jonahtan Vasquez, Richard E. Overstreet, Katherine J. Schultz, Thomas O. Metz, Jessica L. Bade

## Abstract

*Pseudomonas putida* strain KT2440 is a crucial model organism for synthetic biology and bioengineering applications, yet there currently exists no comprehensive metabolomics database comparable to those available for other model organisms. This gap hinders the use of untargeted metabolomics for exploratory analyses in this system. We developed the *P. putida* metabolome reference database (PPMDB v1) to address this limitation by consolidating metabolite information from multiple sources and expanding coverage through computational predictions. The database was constructed by curating metabolites from BioCyc, BiGG, and other literature sources, then computationally expanding this collection using BioTransformer environmental transformation predictions to generate additional predicted metabolites. We enhanced the database’s utility for molecular annotation in metabolomics studies by incorporating analytical properties including collision cross-sections, tandem mass spectra, and gas-phase infrared spectra. These analytical properties were gathered from existing measurement data or predicted using computational tools. We further augmented the database through inclusion of reaction information and pathway annotations, facilitating biological interpretation of metabolomics data. This publicly available resource fills a critical gap in *P. putida* research infrastructure, supporting metabolite annotation and biological interpretation in untargeted metabolomics studies and enabling in-depth exploratory analyses of this important synthetic biology platform at the molecular level.

**GRAPHICAL ABSTRACT:** 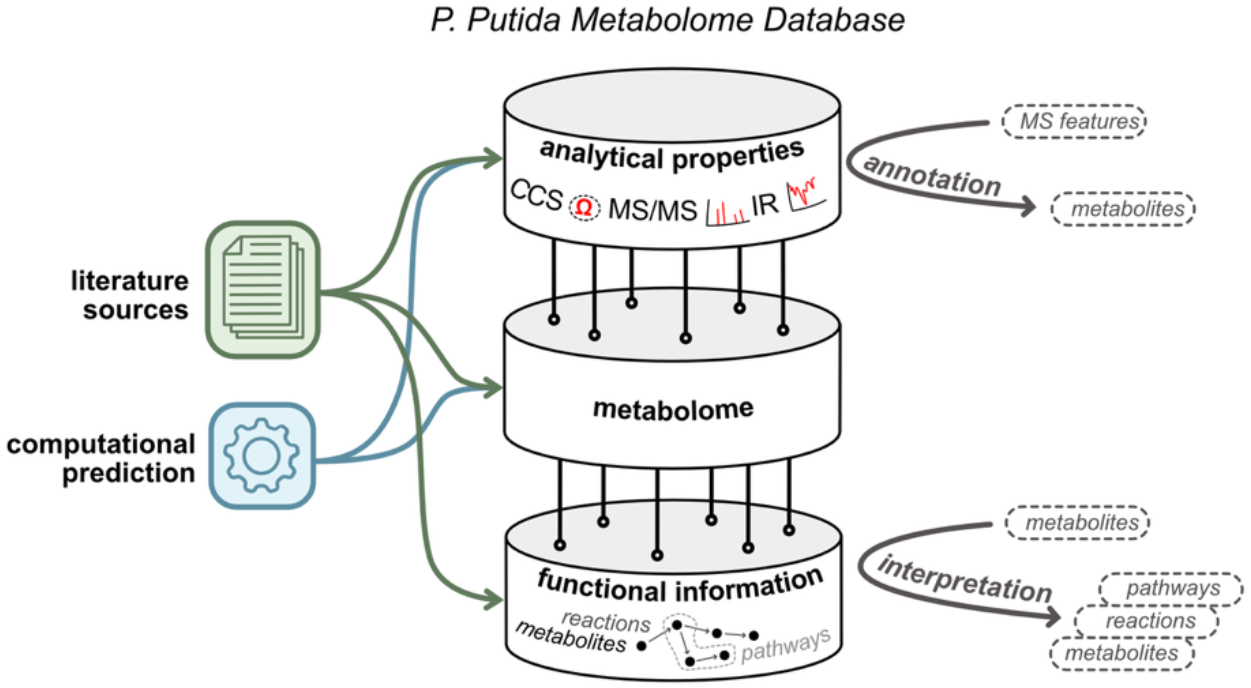

## INTRODUCTION

*Pseudomonas putida*, particularly strain KT2440, has emerged as a cornerstone model in synthetic biology, bioremediation, and bioeconomy applications due to its robust redox capabilities, metabolic adaptability, and resilience under extreme conditions (1). Its role in processes such as biofuel production and chemical valorization underscores its significance as a platform for bioengineering applications. Despite growing interest in harnessing *P. putida*’s metabolic potential, key technical limitations in metabolomics resources hinder exploratory and untargeted analyses of this organism.

Metabolomics seeks to comprehensively profile small-molecule metabolites that drive and are markers for cellular processes, and the inherent proximity of metabolites to phenotype makes it an invaluable tool for probing the molecular underpinnings of complex biological processes (2). Such analyses demand well-characterized metabolite reference data to facilitate the annotation of experimental observations and to contextualize biological insights. Existing databases, especially those for model systems like the Human Metabolome Database (HMDB)(3), *Escherichia coli* Metabolome Database (ECMDB)(4), and Yeast Metabolome Database (YMDB)(5), provide curated metabolite information — including reference data such as mass spectrometry and nuclear magnetic resonance spectra — essential for metabolite annotation and biological hypothesis generation. These tools exemplify the potential of organism-specific metabolome databases to facilitate expanding our understanding of biology in a practical way that supports systems biology and synthetic biology workflows.

No comparable comprehensive metabolomics-focused resource exists for *P. putida*. Existing resources often focus on genomic and pathway reconstructions for KT2440, rather than metabolites, and are limited only to the chemical space of its known central metabolism. This leaves a critical gap in coverage for hundreds of metabolites relevant to engineered strains and niche biotechnological applications and not reflected in *P. putida* central metabolism. Metabolomics researchers are therefore left to rely on disparate resources, such as MassBank of North America (MoNA) [https://mona.fiehnlab.ucdavis.edu], or overly general computational tools with inherent biases that limit their reliability. A centralized, curated reference database is needed to bridge these gaps, integrating experimentally validated and computationally predicted analytical property data to support metabolite annotation needs and downstream biological analyses and interpretation specific to *P. putida*.

In this work, we address these limitations through the creation of the *P. putida* metabolome database (PPMDB, v1), which consolidates observable molecular properties and biological pathway information scattered across literature and experimental studies, and augments existing data using computational predictions. This resource represents a unified network of metabolites, reactions, and pathways aimed at supporting untargeted metabolomics and unlocking the full potential of observations made at the metabolite level. By curating existing data and expanding the chemical space of *P. putida* through environmental transformation predictions, this database establishes a foundation for reliable metabolite annotation and molecular discovery. We make this resource publicly available and encourage community contributions to enhance and broaden its scope.

## MATERIAL AND METHODS

### Initial Curation of Metabolome Compounds

2,078 metabolites were sourced from the BioCyc smart table for all compounds associated with *P. putida* (PPUT160488CYC version 29.0). Manual curation was performed on metadata columns associated with each metabolite. Missing values for structure descriptors (InChI, InChI Key, SMILES) and relevant metadata (molecular weight, monoisotopic mass, formula) were populated by querying PubChem (6) (pubchempy (7), manual search) with the name of the compound. Table entries with duplicate InChI Key values were deduplicated, where applicable, and merged. 1,944 metabolites with necessary metadata are used in the database. 112 entries were missing essential metadata, either from BioCyc, PubChem, or both and were excluded from the final metabolome.

Most of the reaction, gene, and pathway identifiers were sourced using information from the BioCyc SmartTable, where the information was populated from BioCyc’s Pathway/Genome Database (PGDB). While the PGDB was initially generated in 2017 from the *P. putida* KT2440 annotated genome on RefSeq and using the PathoLogic component of Pathway Tools, the PGDB was updated with manual curation in 2019.

Metabolites were additionally sourced from the Biochemical Genetic and Genomic (BiGG) database, which has compartment-specific annotated metabolites for two metabolic reconstructions of *P. putida* KT2440. The iJN746 and iJN1463 (an updated version of the iJN1462) reconstructions are derived from work by Nogales et al. (8,9).

Additional publications were queried using the criteria of articles containing “Pseudomonas putida engineered metabolism”, and “putida” in the title. Only results from peer-reviewed journals published between 2019 (date of last BioCyc curation) and 2024 (date of query) were considered. 58 publications with synthetic biology modifications were found (10-67), which yielded an additional 305 metabolites not already found in the curated metabolic reconstruction databases. Metabolites were manually extracted from the documentation and the previously described methods of querying PubChem were employed to generate metadata.

### Computational Expansion of the Metabolome

An additional 1,965 new compounds were generated through submission of literature compounds through BioTransformer (v3) (68), a software tool that transforms input molecular identifiers according to rules-based reactions that simulate transformations expected in certain sample matrices (e.g., human gut, environment). The ‘Environmental Microbial Transformation’ option was used to predict metabolites arising from environmental microbial degradation of small organic molecules. This collection of reactions is a proxy for predicted breakdown products of xenobiotics and secondary metabolites left in soil or water or exposed to light and represents possible breakdown of metabolites from bacteria, including *P. putida*. A single round (one generation) of transformation was performed, followed by subsequent post-processing and filtering, producing 4,131 predicted metabolites in total.

### Molecular Property Prediction

Mass spectrometry (MS) has emerged as the workhorse technology for measuring metabolomes of biological systems, due to its sensitivity and resolution. To facilitate the adoption and use of the PPMDB in metabolomics studies, we used a suite of computational tools to predict molecular properties for compounds contained within to support annotations from a range of different MS-based analyses. Specifically, we predicted collision cross-sections (CCS), tandem mass spectra (MS/MS), and gas-phase infrared absorption spectra (IR), as these properties can all be measured on MS-based platforms for metabolomics analyses (69-71). CCS values were predicted using the machine learning (ML)-based tools CCSbase (72) and SigmaCCS (73), both of which have test set median relative errors of less than 2%. MS/MS spectra were predicted using QC-GN^2^oMS^2^ (74) and GrAFF-MS (75) using collision energy settings of 10, 20, and 40 eV for both tools; for QC-GN^2^oMS^2^, spectra were predicted for the [M+H]+ adduct, and for GrAFF-MS, spectra were predicted for both [M+H]+ and [M-H]-. For enhanced flexibility in dealing with this collection of prediction tools, each having their own dependencies and system requirements, we used Apptainer to containerize each tool thereby encapsulating the particular environments required for them to run and enabling them to be deployed on a variety of different systems. We additionally implemented standardized interfaces for the containers, making it possible to swap different prediction tools for the same property without any changes to input/output specification in the calling code.IR spectra were predicted using a modified implementation of Graphormer-IR (76,77) capable of making predictions for [M+H]+, [M+Na]+, and [M-H]- adducts. The Graphormer-IR repository was updated to improve its structure, functionality, and performance. Key changes included reorganizing the codebase into a Python package, adding a single-function IR spectra prediction feature, enabling multi-GPU parallel predictions, streamlining dependencies, and replacing command-line arguments with a YAML configuration file for centralized management. The code for the modified implementation can be accessed at https://github.com/pnnl/Graphormer-IR.

### Database Implementation

The metabolome database is implemented in SQLite, with a hierarchical structure to track metabolite-level information, molecular properties, and functional information including metabolic reactions and pathways. The metabolite-level information contained in the database includes primary data about the molecular species that constitute the metabolome, including name, molecular formula, SMILES structure, and metadata on the origin of the compound entry (*i.e*. sourced from literature *vs*. predicted computationally).

Additional metabolite-level metadata like chemical ontological information (at the kingdom, superclass, class, and subclass levels) predicted using ClassyFire (78), alternate names and external IDs (*e.g*. PubChem CID) are also captured. The database also contains molecular properties (mass to charge ratio, *m/z*; collision cross section, CCS; tandem mass spectra, MS/MS; and gas-phase infrared spectra, IR) from both experimental and computational sources to support compound identification from multi-dimensional MS analyses. Functional information in the form of reactions linking metabolites and enzymes are also included in the database. These reactions are further grouped together to form pathways with corresponding metadata. Ultimately this functional information provides a means to connect metabolites with protein-level observations—like transcriptomic or proteomic data—and facilitates biological interpretation by mapping observations onto pathways.

The database was built in an automated fashion from curated data sources using a custom Python build script, which handles database initialization from a schema file and populating the database with data. Metabolite-level data were curated from multiple literature sources and subjected to multiple normalization and cleaning steps for quality control. Molecular property data were sourced from existing values curated from across the literature in previous work (79), or computationally predicted values newly produced in this work. Functional information was curated primarily from BioCyc (80). All code used to build the metabolome database including the database schema definition, and source datasets are available at https://doi.org/10.5281/zenodo.17917884.

## RESULTS

### Assembly of the Metabolome

We initially curated metabolites from BiGG (1,944), BioCyc (1,944), and miscellaneous literature sources (516) (Figure 1). We next expanded the chemical space of the curated metabolites using Biotransformer (68). 2,166 metabolites were used as inputs, resulting in 4,500 predicted metabolites. BiGG and BioCyc produced a highly overlapping set of metabolites (1,820 in common), but both resources provided complementary metadata and reaction information. We merged as much of this complementary information as possible while condensing duplicates to form the final collection of literature-sourced metabolites (2,022). Among the predicted metabolites, some redundancy was observed arising from presence of different charged states (*e.g*. protonated, deprotonated, zwitterionic) coming from the same neutral structure. To mitigate this redundancy, we retained only the neutral structure for each group of compounds without permanent charges. Ultimately, we retained 4,131 predicted metabolites from Biotransformer in our final collection.

**Figure 1.**
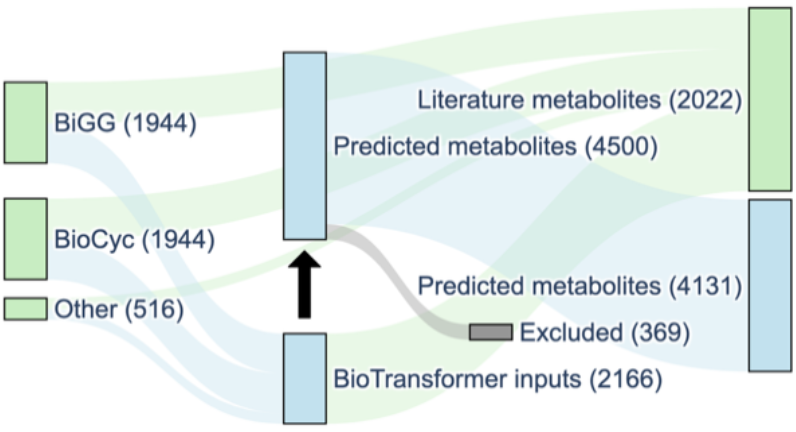
Sankey diagram depicting the initially curated metabolites from primary sources (left), expansion of metabolite chemical space using Biotransformer (center), and final proportions of literature and predicted metabolites in the database (right).

### Chemical Space Coverage

In order to better understand the chemical space spanned by the compounds in the metabolome database, we first performed chemical classification at the levels of kingdom, superclass, class, and subclass using ClassyFire (78) (Figure 2A). All metabolome compounds belong to the “Organic compounds” kingdom, and the primary superclasses with the most representation were “Organic acids and derivatives” (27.5%), “Lipids and lipid-like molecules” (20.8%), “Organoheterocyclic compounds” (15.5%), and “Organic oxygen compounds” (13.6%). The primary constituents of the “Organic acids and derivatives” superclass are the “Carboxylic acids and derivatives” class (61.9%), of which 80.1% come from the “Amino acids, peptides, and analogues” subclass. The “Lipids and lipid-like molecules” superclass is primarily made up of two classes: the “Prenol lipids” (51.3%) and the “Fatty Acyls” (37.6%).

**Figure 2.**
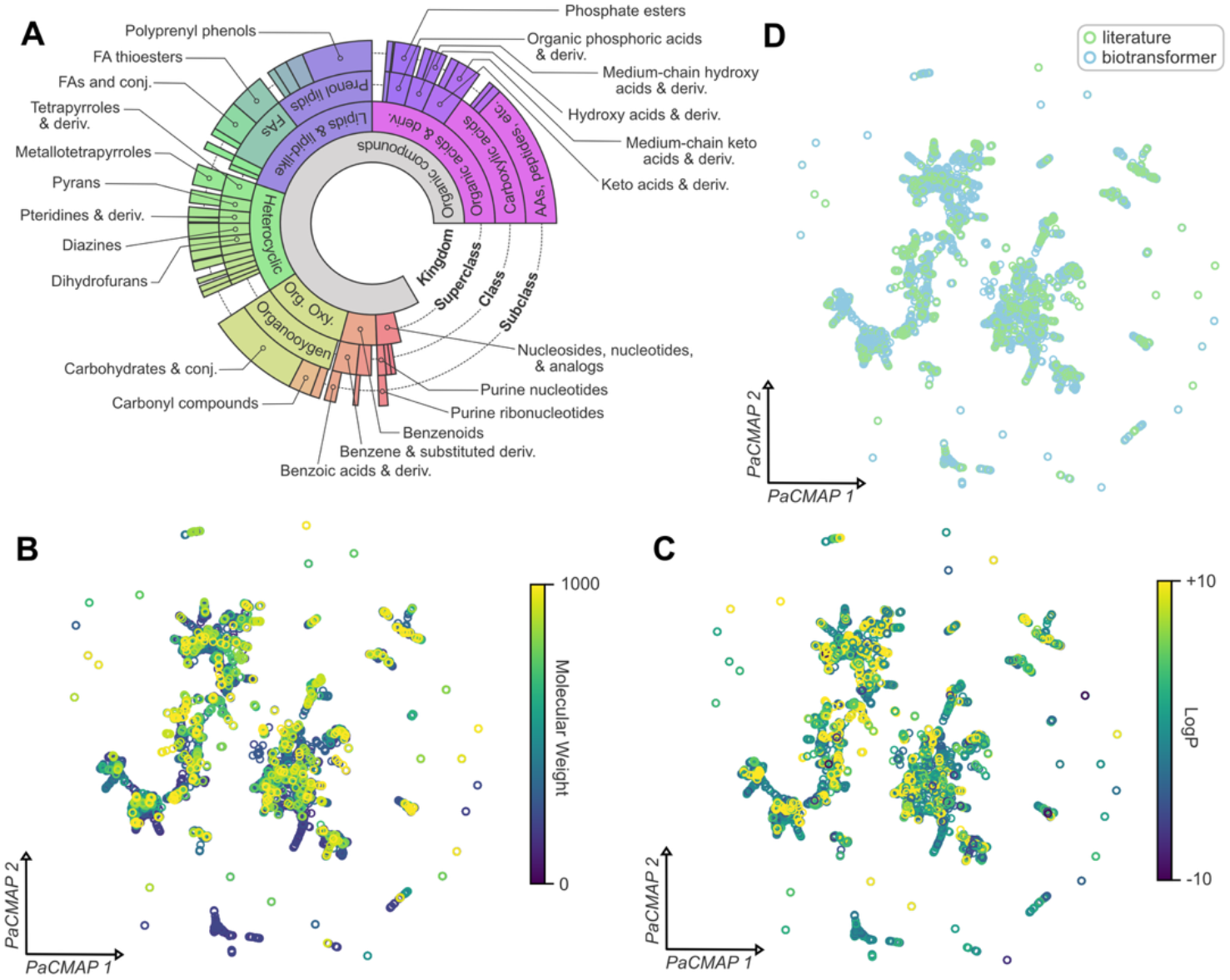
Characterizing the composition and chemical space coverage of the metabolome database. (A) Breakdown of compound classifications at the kingdom, superclass, class, and subclass levels from ClassyFire. (B-D) PacMAP projections of metabolome compounds with points coloured according to compound molecular weight (B), LogP (C), or origin (literature vs. BioTransformer, D)

The main subclass of the “Prenol lipids” is the “Polyprenylphenols” (60.1%) and the two main subclasses of the “Fatty Acyls” are the “Fatty acyl thioesters” (41.8%, mainly Acyl-CoA conjugates) and “Fatty acids and conjugates” (39.1%). The “Organoheterocyclic compounds” superclass was split among several classes and subclasses without a dominant constituent. In contrast, the “Organic oxygen compounds” superclass is composed almost entirely of the “Organooxygen compounds” class (97.9%), which in turn is composed primarily of the “Carbohydrates and carbohydrate conjugates” subclass (73.6%). The classifications above were made based on the complete database—*i.e*., including both literature and predicted metabolites—so we performed an additional analysis with stratification by metabolite origin. We found the general chemical composition between literature and predicted metabolites to be highly similar at the superclass level (literature vs. predicted, respectively: “Organic acids and derivatives” 26.5% vs. 31.0%, “Lipids and lipid-like molecules” 22.1% vs. 23.5%, “Organic oxygen compounds” 19.2% vs. 13.7%, “Organoheterocyclic compounds” 13.4% vs. 18.4%, “Nucleosides, nucleotides, and analogues” 6.3% vs. 2.6%, and “Benzenoids” 5.5% vs. 5.9%), with only marginal differences observed at lower classification levels. Taken together, the chemical classification results demonstrate that the metabolome database represents an overall broad range of chemistries, but certain classes are represented more than others.

We next computed molecular fingerprints and used a pairwise controlled manifold approximation (PaCMAP) (81,82) dimensionality reduction method to produce a two-dimensional projection visually representing the chemical space spanned by the compounds in the metabolome database. For fingerprint generation, we generated 2048-bit Morgan fingerprints for all compounds using RDKit with a connectivity radius of 2; compounds with SMILES where fingerprints could not be generated were discarded (138 SMILES). PaCMAP embeddings were generated using default parameters. Figures 2B-D depict these projections, with compound points colored according to molecular weight (Figure 2B), octanol-water partition coefficient (LogP, Figure 2C), or compound origin (literature vs.

BioTransformer, Figure 2D). Molecular weight and LogP (*i.e*. lipophilicity) are very useful coarse-grained descriptors of the size and composition of the metabolome compounds, and indeed, in the projection plots compounds seem to localize to distinct regions based on these properties. More specifically, the lower molecular weight compounds are more represented near the center of the projected space, with larger compounds distributed more peripherally. Based on LogP, there seems to be an overall trend of increasingly lipophilic compounds in the upper left direction in the projected space. When we compare the chemical space of the original compounds curated from literature sources against those produced computationally using BioTransformer (Figure 2D), we observe that the predicted metabolites do not seem to expand the range of chemical space coverage far beyond the original metabolites but rather increase the depth of chemical space coverage within regions that already contained literature compounds. This observation makes sense given that BioTransformer predicts metabolites from input compounds and there is a limit to the chemical diversity that can be introduced through such a mechanism. These results highlight the value of computational methods for expanding coverage of molecular space, which significantly broaden the aperture of metabolites that can be identified but in a way that maintains the general landscape of the chemical space.

### Analytical Properties

A key function of PPMDB is to support annotation of compounds that are observed experimentally using various MS-based analytical methods. We therefore include relevant analytical properties (*m/z*, CCS, MS/MS spectra, and gas-phase IR spectra) that would support compound annotation from a variety of MS-based metabolomics analyses ranging from routine to highly specialized. Existing values for these properties were previously curated from a variety of published sources (79), and in addition to these we performed computational predictions to further expand the coverage, using a modular workflow with containerized prediction tools. Figure 3A depicts the molecular property coverage of the database, broken down by compound source (“literature” or “biotransformer”) and adduct (“[M+H]+”, “[M+Na]+”, or “[M-H]-”). The molecular property coverage for all properties is highest among the compounds that are derived from literature sources. Among the literature compounds, the [M+Na]+ adduct has very low MS/MS coverage, which is attributable to a limitation of the computational tools used to predict these spectra, which predict either protonated adducts exclusively (QC-GN^2^oMS^2^ (74)) or protonated and deprotonated adducts exclusively (GrAFF-MS (75)). Promisingly, among the literature compounds that have associated molecular properties, there is a high degree of overlapping coverage across all three molecular properties (Figure 3A, Venn diagrams).

**Figure 3.**
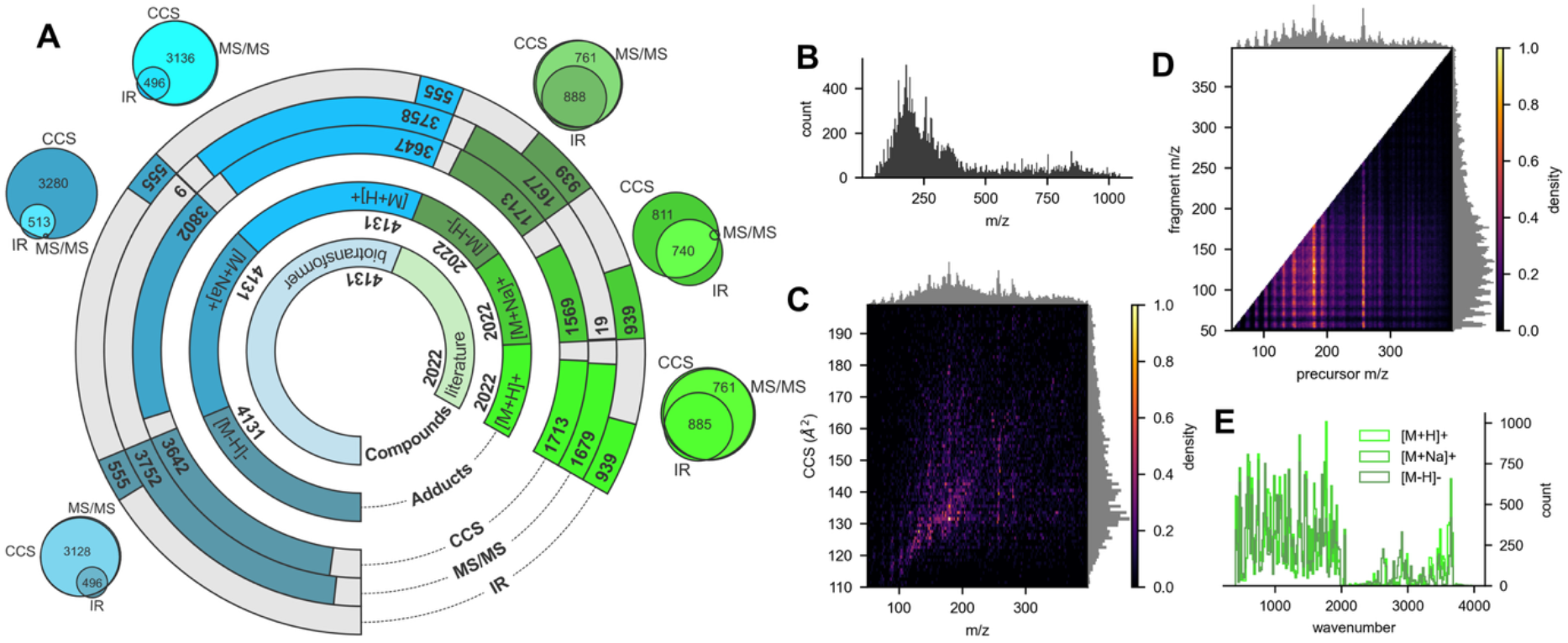
Characterizing the analytical property coverage for compounds in the metabolome database. (A) Annular plot showing molecular property coverage for metabolome compounds, broken down by origin (literature vs. BioTransformer) and adduct ([M+H]+, [M+Na]+, and [M-H]-). Additional Venn diagrams are included outside of the annular plot which depict the molecular property coverage in terms of combinations of the properties. Counts are only specified for significantly overlapping portions (B) Overall distribution of precursor m/z values in the database. (C) Two-dimensional histogram showing the distribution of m/z and CCS values in the database. Grey histograms above and to the right are the corresponding single-dimension histograms for m/z and CCS. (D) Two-dimensional histogram depicting the characteristics of the MS/MS spectra contained in the database. In this plot, the precursor m/z is represented on the x-axis and fragment m/z from the MS/MS spectra are represented on the y-axis. Vertical bands in this plot are indicative of precursors that produce a variety of fragments across a wide m/z range, while horizontal bands suggest fragments that are common to multiple precursors. Grey histograms above and to the right are corresponding single-dimension histograms for precursor and fragment m/z. (E) Histograms of IR peaks from all gas phase IR spectra contained in the database, broken down by adduct.

We next examined the distributions of molecular property values contained in the database to understand the range of identifiable chemical space in terms of analytical measurements. Figure 3B depicts the distribution of precursor *m/z* for all adducts in the database, from which it is clear the metabolome is dominated by small molecules in the 50-400 *m/z* range. Considering that IMS is typically coupled with MS, it makes sense to characterize the CCS coverage along with precursor *m/z* (Figure 3C). From this representation of the IMS-MS conformational space (83), we see the expected correlation between precursor *m/z* and CCS with deviations from the main trend demonstrating coverage across multiple structurally distinct chemical classes. Figure 3D depicts the extent of the MS/MS spectral information present in the database, in terms of precursor and fragment *m/z*, and Figure 3E depicts the overall information content of the predicted IR spectra in the database in terms of wavenumber, broken down by adduct type. While there are some subtle differences in the absorbance bands represented among the IR spectra from different adducts, at a high level each captures roughly similar range and proportion of absorbance bands (representing different functional groups).

### Functional Information

Functional analyses, which connect metabolite identities and abundances with enzymes and metabolic pathways, represent a critical link between changes at the metabolome level and actionable biological hypotheses. To this end, we have included functional information that links metabolites, enzymes and pathways (curated from KEGG(84) and BioCyc(80)) within the metabolome database. We also include identifiers from KEGG and BioCyc to enable cross-referencing with these external resources. In total, 972 metabolites take part in 1050 reactions that are associated with 944 pathways. The top 20 pathways from KEGG and BioCyc (excluding overview pathways) according to the number of metabolites they include are summarized in Figures 4A and 4B, respectively. These top pathways span many areas of cellular metabolism from processing of amino acids, lipids, and small molecules to biosynthesis of critical cofactors and energy production. The top enzymes present in the database according to the number of metabolites they are associated with (Figure 4C) include those involved in Fe-S cluster system, lipid metabolism, and cofactor biosynthesis. Having this rich functional information integrated within the metabolome database enhances the impact of metabolomic studies by enabling the interpretation of important metabolites within a broader biological context, without the need to separately seek out biological connections from external resources.

**Figure 4.**
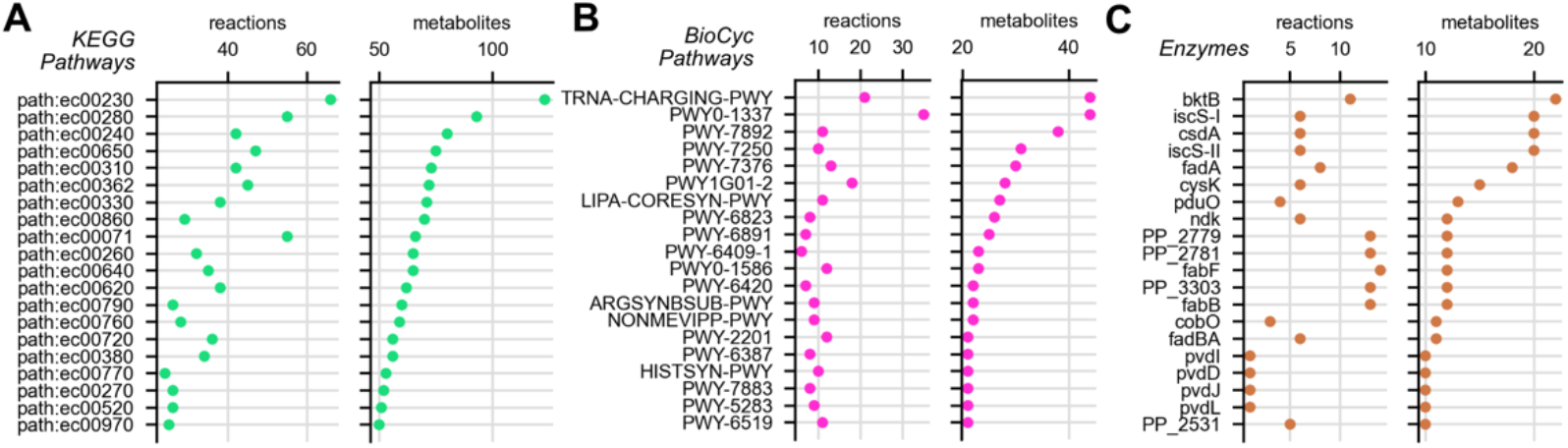
Summaries of top pathways and enzymes in the database by associated reaction and metabolite counts. Top 20 pathways (A: KEGG, B: BioCyc) according to count of associated metabolites (right panels) as well as counts of associated reactions (left panels). (C) Top 20 enzymes according to count of associated metabolites (right panel) with counts of associated reactions (left panel).

## DISCUSSION

The *P. putida* metabolome database presented in this work represents a valuable and novel resource supporting metabolomic profiling and biological contextualization of metabolites towards bioengineering applications in a way not previously possible. Beyond cataloguing metabolites that define the chemical landscape of *P. putida*, the database directly supports practical metabolomics data analysis objectives including metabolite annotation and biological hypothesis generation through functional analyses. The PPMDB consolidates information curated from multiple sources, creating a centralized resource that eliminates the need to search across disparate platforms and ensuring high-quality, reliable data for robust and accurate results.

Previous work in this area has focused on genome-scale models of *P. putida* metabolism, such as the iJN1462 (9) model of metabolism and the more recent iPpu1676-ME (85) model of metabolism and gene expression, which mechanistically integrate gene expression and metabolism to optimize resource allocation for growth. These genome-scale models excel in predicting and optimizing macro-level phenotypes, such as biomass production, substrate consumption, and targeted molecule production. However, their gene-centric construction and focus on transcriptional and proteomic readouts mean they are less suited for untargeted or exploratory analyses, particularly metabolomics. In contrast, the metabolome database presented in this work is developed specifically to support untargeted and exploratory metabolomics analyses, which synergizes perfectly with existing resources that support macro-level phenotypic analysis objectives. In summary, we believe this resource will meaningfully expand the informatic toolkit for interrogation of *P. putida* at the molecular level.

## AUTHOR CONTRIBUTIONS

DHR: Data curation, Formal analysis, Visualization, Software, Supervision, Writing – original draft. CHC: Data curation, Software, Writing – original draft. JV: Data curation. REO: Data curation, Software. KJS: Conceptualization, Funding acquisition. TOM: Conceptualization, Funding acquisition, Project administration, Supervision. JLB: Data curation, Methodology, Writing – original draft. All authors: Writing – review & editing.

## ACKNOWLEDGEMENTS

This work was supported by the Pacific Northwest National Laboratory (PNNL) Laboratory Directed Research and Development program under the Predictive Phenomics Initiative. Predictions were performed using PNNL Institutional Computing and high-performance computing resources of the Environmental Molecular Sciences Laboratory, which is a U.S. Department of Energy (DOE) Office of Science User Facility. PNNL is a multiprogram national laboratory operated for the DOE by Battelle Memorial Institute under DOE Contract No. DE-AC05-76RL01830. PNNL-SA-218685.

## DATA AVAILABILITY

Database files and source datasets are available on Zenodo at https://doi.org/10.5281/zenodo.17917884.

